# An enzyme-free, cold-process acoustic method for gentle and effective tissue dissociation

**DOI:** 10.1101/2023.10.03.560704

**Authors:** Melanie A. MacMullan, Marlee Busalacchi, Sophie Quisling, Brian Quast, Johnna Pullen, Sydney Addington, Vibhu Vivek, Steve Levers, Kristin Renkema

**Author notes:** Corresponding authors, (M.M.), (K.R.). designates first author (Melanie A. MacMullan, Marlee Busalacchi).

## Abstract

As biological advances continue to improve the resolution of genomic and proteomic studies, the quality of single cell suspensions is becoming increasingly important. While conventional approaches use enzymes which may require heat Abbreviations: Bulk Lateral Ultrasound (BLU) activation to break down extracellular tissue matrices and gain access to single cells, recent studies have suggested that these harsh biochemical and heat-based treatments may result in genomic and proteomic modulation. To minimize these dissociation artifacts, we have developed an instrument for dissociating cells from various tissue matrices using Bulk Lateral Ultrasound (BLU™) energy. This enzyme-free, gentle mechanical dissociation maintains sample temperatures below 8°C for the duration of processing, resulting in high-fidelity single cell suspensions with comparable viability and live cell counts to those obtained with conventional enzymatic dissociations. Here, in murine-derived brain, heart, lung, and B16 melanoma tumor tissue dissociated by either BLU or by a commercially available dissociation kit which uses enzymes and heat, we compare cell viability and expression of population-specific immunological markers. The dramatic differences observed in cell surface expression suggest that cells dissociated using enzymes and heat may be experiencing stress-induced changes post-harvest that could impact conclusions and impede research progress. Alternatively, using gentle mechanical dissociation with BLU, we demonstrate the preservation of these markers and enable a minimally invasive alternative to obtaining high integrity single cell suspensions.

**Highlights:**

- Novel, acoustic energy-based, enzyme-free dissociation improves single cell suspensions
- Enzymatic dissociation diminishes cell counts, viability, and surface marker expression
- Immunophenotyping reveals marker preservation by acoustic-based dissociaton

## 1. Introduction

Proper dissociation of tissue samples harvested from patients and research animals is essential for a growing number of downstream processes including fluorescence-activated cell sorting (FACS), single cell RNA sequencing (scRNA-seq), and single cell proteomics. Conventional methods for dissociation, which typically involve the use of various enzymes to detach cells from one another and from their extracellular matrix, result in poorly reproducible single cell suspensions that are often low in viability (1). In addition to the inconsistencies in sample processing that arise from tissue dissociation with enzymes, higher resolution analyses are revealing artificial changes in genomic and proteomic signals that result from these biochemical treatments (2,3).

Many enzymes require incubation at 37°C for activation, introducing thermal stresses that may cause cells to deviate from their original states. Even during short periods of heat shock, mammalian cells illicit an orchestrated stress response resulting in largely modulated transcriptomes (4). Significant transcriptomic changes have also been observed by O’Flanagan et al. in an scRNA-seq analysis of cancerous tissues and cell lines dissociated with enzymes at 37°C when compared to a cold dissociation protocol (5). Other studies have similarly shown cell population changes to microglia in the brain and gene expression bias in skeletal myoblasts which increases with enzymatic digestion time (6,7). Although cold shock of specimens has also been demonstrated to impact cellular integrity, the majority of tissue samples isolated clinically are not able to be analyzed immediately and must be placed on ice for transportation to preserve cell viability post-harvest (8,9). While the influence of an initial cold shock may be unavoidable, these studies demonstrate that subjecting tissue samples to further heating and cooling cycles are likely substantially altering single cell suspensions post-harvest, leading to questions about the integrity of downstream genomic and proteomic data.

The heterogeneity of the immune response in both healthy and diseased tissues can reveal novel insights both into physiological homeostatic function and disease (10). Immune cells occupy a range of sizes and embody a variety of functions which work together in a delicate balance, highlighting the importance of identifying both common and rare populations. Cell surface markers unique to immune cell subpopulations are typically used to identify cellular subtypes isolated from various tissues and tumor microenvironments (11). Flow cytometry-based cell analyses are commonly used to assess relative quantities of immunological subpopulations and sorted cells are often used in downstream analyses. For both methods, single cell suspensions are required as input material for proper staining and analysis and to prevent instrument clogging. While enzyme and heat-based dissociation protocols have often preceded flow cytometry and FACS, recent studies have demonstrated a significant loss of certain immunological cell markers following dissociation with enzymes (12,13). Further, single-cell analysis methods often target rare cell types, making the accurate preservation of varied cell populations and cell markers essential for these applications.

As an alternative to enzyme and heat-based dissociation, mechanical methods of dissociation have been attempted with limited success. Unfortunately, many of these methods have the disadvantage of leaving cells trapped in extracellular matrices, resulting in a low number of viable cells dissociated from various tissue sources (14). Our recently developed SimpleFlow™ technology employs Bulk Lateral Ultrasound (BLU) energy for an effective but gentle tissue dissociation that maintains samples below 8°C in a temperature-controlled chamber. To compare dissociation of various murine tissue types by BLU to dissociation by enzymes and heat, we used commercially available enzyme and heat-based dissociation kits from Miltenyi (Miltenyi Biotec, Charlestown, MA, USA). By flow cytometry-based cell analysis, we evaluated viable cell numbers and the relative expression of several population-differentiating immunological markers on tissue samples harvested from commercially purchased mice, a population of which were injected with B16 melanoma tumors. All mice had previously been exposed to a variety of microbes in order to elicit immune responses (manuscript in prep, K. Renkema). By characterizing the expression of these markers in cells dissociated from equivalent tissue samples by either method, we demonstrate that several markers known to be impacted by conventional dissociation methods are preserved with BLU dissociation and identify several other immunological markers of interest lost by enzymatic dissociation.

## 2. Methods

### 2.1. Animal collection

Female C57Bl/6 mice were purchased from Charles River Laboratories (Mattawan, MI, USA) and experiments were conducted when the mice were 8-12 weeks old. Female pet store mice were purchased from a local pet store. All mice were exposed to multiple microbes in order to measure immune responses to infection. All animal experiments were done with approved Institutional Animal Care and Use Committee protocols at Grand Valley State University (#23-08-A).

### 2.2. B16 Melanoma injection

To obtain tumor tissue, B16-F10 cells were purchased from American Type Culture Collection (ATCC) and cultured to ∼80% confluency, counted, and prepared in sterile PBS for injection. Briefly, 2×10^5^ B16-F10 cells in sterile PBS were subcutaneously injected into shaved bilateral flanks of C57Bl/6 mice, as previously described (15,16). Twelve days post-injection, tumors were isolated and prepared as described in section 2.4.1.

### 2.3. Murine tissue harvest

To compare BLU dissociation to more conventional dissociation techniques using enzymes and heat, various murine tissues were harvested from female C57Bl/6 or pet store mice. Whole brains and hearts, pairs of lungs, and B16 melanoma tumor tissue were used in these experiments. Twelve days post B16 melanoma injection, C57Bl/6 mice were euthanized and bilateral tumors from the flanks of each mouse, as well as whole brains, were harvested. Whole hearts and pairs of lungs were harvested from pet store mice.

### 2.4. Tissue dissociation

#### 2.4.1. B16 Melanoma tumor dissociation

B16 melanoma tumors (N=6) were obtained from the bilateral flanks of three C57BL/6 mice 12 days post tumor injection and divided into ten total samples. One sample from each tumor was used to examine two dissociation conditions: BLU energy and a commercially available enzyme and heat dissociation kit, the Miltenyi Tumor Dissociation Kit for mice (Cat. No. 130-096-730). Fat, fibrous and necrotic areas were removed from the tumors which were subsequently divided into aliquots of relatively equal mass, approximately 100-120 mg.

For samples dissociated by BLU (N=5), tumor tissue and 250 µL of RPMI were added to the mincing chamber of the SimpleFlow cartridge and minced manually before being transferred to the dissociation chamber. An additional 500 µL of RPMI was added to the mincing chamber incrementally to rinse the chamber and the mincer. Rinsed sample was added to the dissociation chamber with the remainder of the sample. The sample was then dissociated using BLU energy for two minutes while maintaining temperatures below 8°C.

According to the Miltenyi Tumor Dissociation Kit protocol, five tumor tissue samples were cut into 2-4 mm pieces and then transferred to a gentleMACS™ C tube (Cat. No. 130-093-237) containing a 162.5 µL cocktail of three enzymes provided in the kit. Samples were dissociated on the Miltenyi gentleMACS Octo Dissociator (Cat. No. 130-096-427) using the 15-minute 37C_m_TDK_1 program.

#### 2.4.2. Brain tissue dissociation

Whole brains were obtained from three C57BL/6 mice 12 days post B16 melanoma injection. Brains were washed with cold D-PBS and excess spinal cord and other tissue was removed. Brain tissue was divided into ten total samples with similar weights (∼100-120 mg). Two conditions were examined with five samples (N=5) each: BLU energy and a combination of enzymes and heat using the Miltenyi Adult Brain Dissociation Kit for mouse and rat (Cat. No. 130-107-677).

For the samples dissociated by BLU, one of five aliquots of brain tissue and 250 µL of RPMI was added to each SimpleFlow cartridge mincing chamber and minced manually. Minced brain tissue was then transferred to the dissociation chamber. An additional 500 µL of RPMI was added incrementally to rinse the mincing chamber and was then transferred to the dissociation chamber with the remaining sample. Cartridges were stored on ice until processing on SimpleFlow, where the three samples were dissociated for 2 minutes using BLU energy while maintaining temperatures below 8°C.

According to the Miltenyi Adult Brain Dissociation Kit for mouse and rat protocol, the five samples for enzyme and heat dissociation were cut into smaller pieces before being transferred to a gentleMACS C tube. Two enzyme mixtures were prepared by combining one enzyme and one buffer each. Enzyme Mix 1 was added to a fresh gentleMACS C tube, followed by the tissue sample, and then by Enzyme Mix 2. Samples were dissociated on the gentleMACS Octo Dissociator using the 28-minute 37C_ABDK_01 program.

#### 2.4.3. Heart tissue dissociation

Heart tissue was obtained from ten pet store mice and divided into ten total samples with similar weights. Five samples each (*N* = 5) were used to examine two conditions: BLU energy and the enzyme and heat based Miltenyi adult mouse heart dissociation protocol using Miltenyi’s Multi Tissue Dissociation Kit 2 (Cat. No. 130-110-203). Hearts were removed and tissue was washed in cold D-PBS to rinse extra blood from the tissue. Excess tissue was then removed from the hearts. Heart tissue was divided into ten total samples with similar weights.

For samples dissociated by BLU energy, heart tissue and 250 µL of RPMI was added to the SimpleFlow cartridge mincing chamber and minced manually before being transferred to the dissociation chamber. An additional 500 µL of RPMI was added incrementally to the mincing chamber to rinse the cartridge and the mincing blade. The rinse was transferred into the dissociation chamber with the remainder of the sample. Cartridges were stored on ice until processing with BLU energy for 2 minutes while maintaining temperatures below 8°C.

According to the Miltenyi heart dissociation protocol, a 2.5 mL cocktail of three enzymes was placed into a gentleMACS C tube before adding tissue. Samples were dissociated on the gentleMACS Octo Dissociator using the 57-minute 37C_Multi_G program. After the incubation program was complete, an additional 7.5 mL of RPMI-20% FBS was added to the sample tubes.

#### 2.4.4. Lung tissue dissociation

Pairs of lungs were obtained from ten pet store mice and divided into ten total samples of similar weights. Five samples each were used to examine two conditions: a two-phase BLU energy protocol, and the enzyme and heat based Miltenyi Lung Dissociation Kit for mice (Cat. No. 130-095-927). Excess tissue and blood were removed from the lungs in petri dishes containing cold PBS. Pairs of lungs were separated into individual lungs before being divided into ten total samples. Lungs samples were dried and weighed prior to dissociation. Each lung was cut into smaller pieces prior to dissociation.

For the samples dissociated by a two-phase BLU energy protocol, lung tissue and 250 µL of RPMI were added into a SimpleFlow cartridge mincing chamber and minced manually. Minced lung tissue was then transferred to the dissociation chamber. An additional 500 µL of RPMI was added to rinse the mincing chamber incrementally and was then transferred to the dissociation chamber. Cartridges were stored on ice until processing on SimpleFlow, where the five samples experienced high BLU energy for 1 minute followed by low BLU energy for 1 minute. Both protocols were performed while maintaining temperatures below 8°C.

According to the manufacturer’s protocol, the remaining tissue samples (*N* = 5) were transferred to a gentleMACS C tube containing a 2.515 mL cocktail of two enzymes. Samples were dissociated on the gentleMACS Octo Dissociator using the 31-minute 37C_m_LDK_1 program.

### 2.5. Gravity Filtration

For all samples, the entire volume was removed by pipette from the SimpleFlow cartridge dissociation chamber or gentleMACS C tube and placed onto a 70-micron filter in a 50 mL conical tube. 50 mL tubes were stored on ice during the gravity filtration process. The cartridge or gentleMACS C tube was rinsed with cold RPMI or RPMI-2%. The rinsed volume was transferred onto the 70-micron filter and a syringe plunger was used in a grinding motion on the filter to pass residual tissue.

To maintain consistency, centrifugation settings from Miltenyi kit protocols were applied to single cell suspensions from both dissociation methods. Single cell suspensions from tumors were centrifuged for 5 minutes at 300 xg and 4°C. Single cell suspensions from brain and lung tissue were centrifuged for 10 minutes at 300 xg and 4°C. Single cell suspensions from heart tissue was centrifuged for 5 minutes at 600 xg and 4°C. The supernatant was completely removed from all samples. Cells from tumor and lung samples were resuspended in RPMI, and cells from heart and brain samples isolated by enzymatic and heat dissociation methods underwent debris removal and red blood cell (RBC) lysis. These steps were not applied to BLU dissociated suspensions.

### 2.6. Debris Removal

For single cell suspensions enzymatically dissociated from heart and brain tissue, the cell pellet was resuspended in PBS and cold Debris Removal Solution (Miltenyi Cat. No. 130-109-398) was added. The samples were mixed well using a pipette. Additional PBS was gently overlayed. All samples were centrifuged for 10 minutes at 3000 xg and 4°C, creating three phases. The top two phases were aspirated and discarded. More PBS was added and mixed gently by inverting. Samples were centrifuged for 10 minutes at 1000 xg and 4°C. The supernatant was aspirated completely, and cells were resuspended in for RBC lysis.

### 2.7. RBC Lysis

Single cell suspensions enzymatically dissociated from brain and heart tissue were resuspended in ammonium-chloride-potassium (ACK) lysing buffer (Miltenyi Red Blood Cell Lysis Solution, Cat. No. 130-094-183) and incubated for 2 minutes at room temperature. For brain cell suspensions, the cell-buffer solution was then diluted in FACS Buffer (PBS with 2% FBS) and centrifuged for 10 minutes at 300 xg and 4°C. The supernatant was aspirated completely, and cells were resuspended in RPMI-2%. Heart cell suspensions were loaded directly into the centrifuge without dilution and centrifuged for 5 minutes at 600 xg and 4°C. Supernatant was aspirated completely and cells were washed once with PBS and centrifuged for 5 minutes at 600 xg and 4°C. The supernatant was again aspirated completely, and cells were resuspended in FACS buffer.

### 2.8. Antibody preparation and immunostaining

Single cell suspensions were aliquoted into a 96-well plate. Plates were centrifuged for 5 minutes at 300 xg and 4°C and the supernatant was aspirated. Samples were then incubated with an Fc block (BioLegend, Cat. No. 156604) for 10 minutes at 4°C. FACS buffer was added to each sample, the plate was centrifuged for 5 minutes at 300 xg and 4°C, and the supernatant was aspirated. Next extracellular staining antibodies were added, and the samples were incubated for 20-30 minutes at 4°C while protected from light. Antibodies were combined in different panels comprising of an appropriate selection from the following: a 1:400 dilution of Alexa Fluor™ 750 anti-murine CD8 (BioLegend, Cat. No. 100766), a 1:400 dilution of Brilliant Violet™ 605 anti-murine CD4 (BioLegend, Cat. No. 100548), a 1:400 dilution of PerCP-Cyanine5.5 anti-murine CD3 (BioLegend, Cat. No. 100217), a 1:500 dilution of FITC anti-murine CD45 (BioLegend, Cat. No. 103107), a 1:400 dilution of APC/Fire™ 750 anti-murine CD115 (BioLegend, Cat. No. 135536), a 1:400 dilution of Brilliant Violet™ 421 anti-murine CD11b (BioLegend, Cat. No. 101251), a 1:400 dilution of Brilliant Violet™ 510 anti-murine Ly6C (BioLegend, Cat. No. 128033), a 1:400 dilution of Brilliant Violet™ 605 anti-murine MHC-II (BioLegend, Cat. No. 107639), a 1:400 dilution of PE-Cyanine7 anti-murine F4/80 (BioLegend, Cat. No. 139314), a 1:500 dilution of Alexa Fluor™ 700 anti-murine CD86 (BioLegend, Cat. No. 123130), a 1:500 dilution of Alexa Fluor™ 647 anti-mouse CD206 (BioLegend, Cat. No. 151508), a 1:400 dilution of APC/Fire™ 750 anti-murine CD11c (BioLegend, Cat. No. 117351), a 1:400 dilution of PE-Cyanine7 anti-murine CD144 (BioLegend, Cat. No. 138015), a 1:400 dilution of PerCP-Cyanine 5.5 anti-murine CD140a (BioLegend, Cat. No. 135913), a 1:400 dilution of PE-Cyanine7 anti-murine CD64 (BioLegend, Cat. No. 161007), a 1:500 dilution of Alexa Fluor™ 700 anti-murine CD24 (BioLegend, Cat. No. 101835), and a 1:500 dilution of Alexa Fluor™ 647 anti-murine Siglec-F (BioLegend, Cat. No. 155519). Depending on the fluorophores selected in each panel, either the LIVE/DEAD™ Fixable Violet Dead Cell Stain Kit (Invitrogen, Cat. No. L34955) or the LIVE/DEAD™ Fixable Red Dead Cell Stain Kit (Invitrogen, Cat. No. L34971) were included in the panel according to the manufacturer’s protocol. All antibody dilutions were made in FACS buffer. Following staining, FACS buffer was added to wash each sample, and the plate was centrifuged at 300 xg and 4°C. The supernatant was removed.

### 2.9. Fixation and Permeabilization

1X Fix Concentrate from the True-Nuclear™ Transcription Factor Fixation/Permeabilization Buffer Set (BioLegend, Cat. No. 424401) was added to each sample, and then incubated for 30 minutes at 4°C, protected from light. Cells were then washed three times in the 1X Permeabilization Buffer from the kit with centrifugation for 5 minutes at 300 xg and 4°C followed by complete aspiration of the supernatant. Cells were then resuspended in a 1:10 dilution of ThermoFisher Scientific CountBright™ beads (Cat. No C36950) in FACS buffer, transferred to a FACS tube, briefly vortexed, and analyzed on the Beckman Coulter Cytoflex.

### 2.10. Data Analysis

FlowJo v10.9.0 was used to determine cell counts, discriminate live cell populations, and quantify cell surface expression by median fluorescent intensity (MFI). GraphPad Prism v10 was used to generate plots and run statistical analyses. Unpaired two-tailed t-tests were used to evaluate statistical significance.

## 3. Results

### 3.1. Dissociation by BLU energy yields healthy single cell suspensions

All data presented here are representative of five biological replicates of cells dissociated from mouse tissue by SimpleFlow Bulk Lateral Ultrasound (BLU) energy or by tissue-specific Miltenyi dissociation kits (Fig. 1A). Aside from analyses in lung tissue, the results presented here were observed in previous comparisons, though only data from one experiment are shown here. Tissue samples were processed to minimize variability in starting material and expression (Supp. Table 1). The initial metrics used to evaluate the success of tissue dissociation were the viability of the dissociated cells and the number of live cells dissociated per milligram of input tissue. We have occasionally observed, in work that has not been peer-reviewed, that viability post BLU energy dissociation is artificially reduced due to the absence of enzymatic or other chemical methods for dead cell removal (17–20). In this study, differences in viability across tissue types, excluding tumor, were nonsignificant (Fig. 1B). Further, the number of dissociated live cells per milligram of tissue by BLU are consistent with those dissociated by enzymes and heat for all tissue types (excluding lung; Fig. 1C).

**Figure 1.**
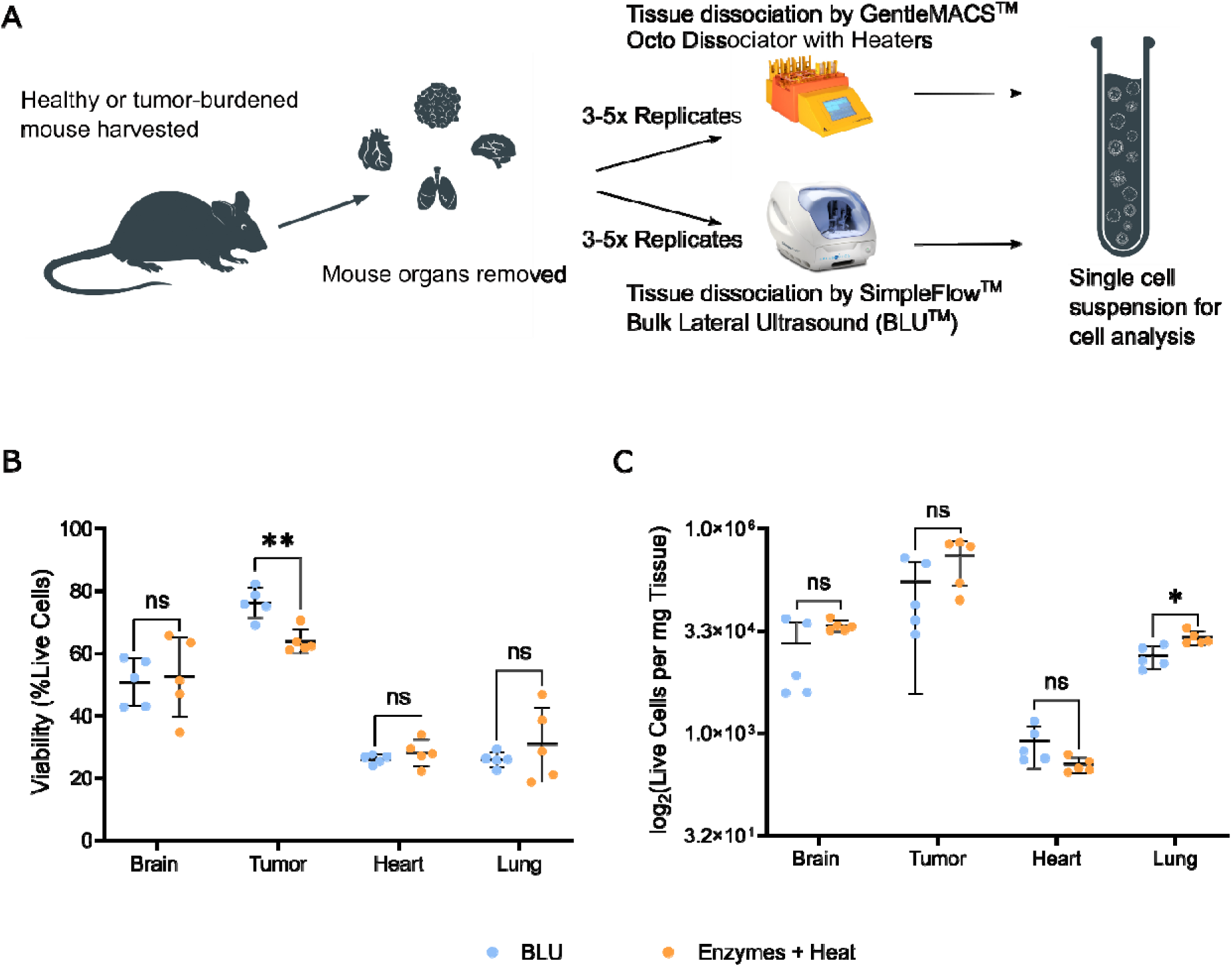
Experimental methods for comparing murine tissue dissociation by a conventional enzymatic approach and a novel mechanical approach. (A) Various organs harvested from mice were aliquoted to obtain samples of equal masses for processing with enzymes by GentleMACS™ Octo Dissociator with Heaters or by SimpleFlow™ BLU™ energy. (B) Viability and (C) cell counts of resulting single cell suspensions were compared between dissociation conditions. ns = not significant, * = p-value ≤ 0.05, ** = p-value ≤ 0.01.

Cells dissociated from the brain had an average viability of 51.5% by BLU and 56.5% by enzymes and heat. From the tumor, heart, and lung, cells were 74.0%, 36.7%, and 26.0% viable respectively when dissociated by BLU and 56.2%, 33.9%, and 30.8% by enzymes and heat. The number of live cells per milligram of brain, tumor, heart, and lung tissue ranged from 1.4×10^4^, 2.0×10^5^, 3.6×10^4^, and 1.4×10^4^ by BLU to 2.6×10^4^, 3.3×10^5^, 1.0×10^4^, and 2.6×10^4^ by enzymes and heat. Based on unpaired two-tailed t-tests, viability and live cell counts were not significantly different by either dissociation method for most tissue types, indicating that BLU dissociation yields a sufficiently healthy single cell suspension for downstream analyses including cell surface marker expression evaluation by flow cytometry.

### 3.2. Tissue dissociation by BLU resulted in a higher yield of CD8+ T Cells

Cells from each tissue type dissociated by either method were evaluated for expression of population-specific immune cell markers. A representative gating strategy from heart tissue is shown in Fig. 2A, where singlet cells were initially selected and a viability dye was used to exclude dead cells from further analysis. Those cells were then analyzed for expression of common helper T cell and cytotoxic T cell markers, CD4 and CD8, respectively. Populations which fell in the first quadrant (Q1) were considered CD4+CD8-, those in Q2 were considered CD4+CD8+, those in Q3 were considered CD4-CD8+ and those in Q4 were considered CD4-CD8-. Further studies compared the median fluorescent intensity (MFI) of CD3 (Fig. 2B, D) and CD8 (Fig. 2C, E) on cells dissociated from all tissue types by both BLU and the enzyme and heat approach as well as the number of CD8+ T cells dissociated per milligram of tissue (Fig. 2F).

**Figure 2.**
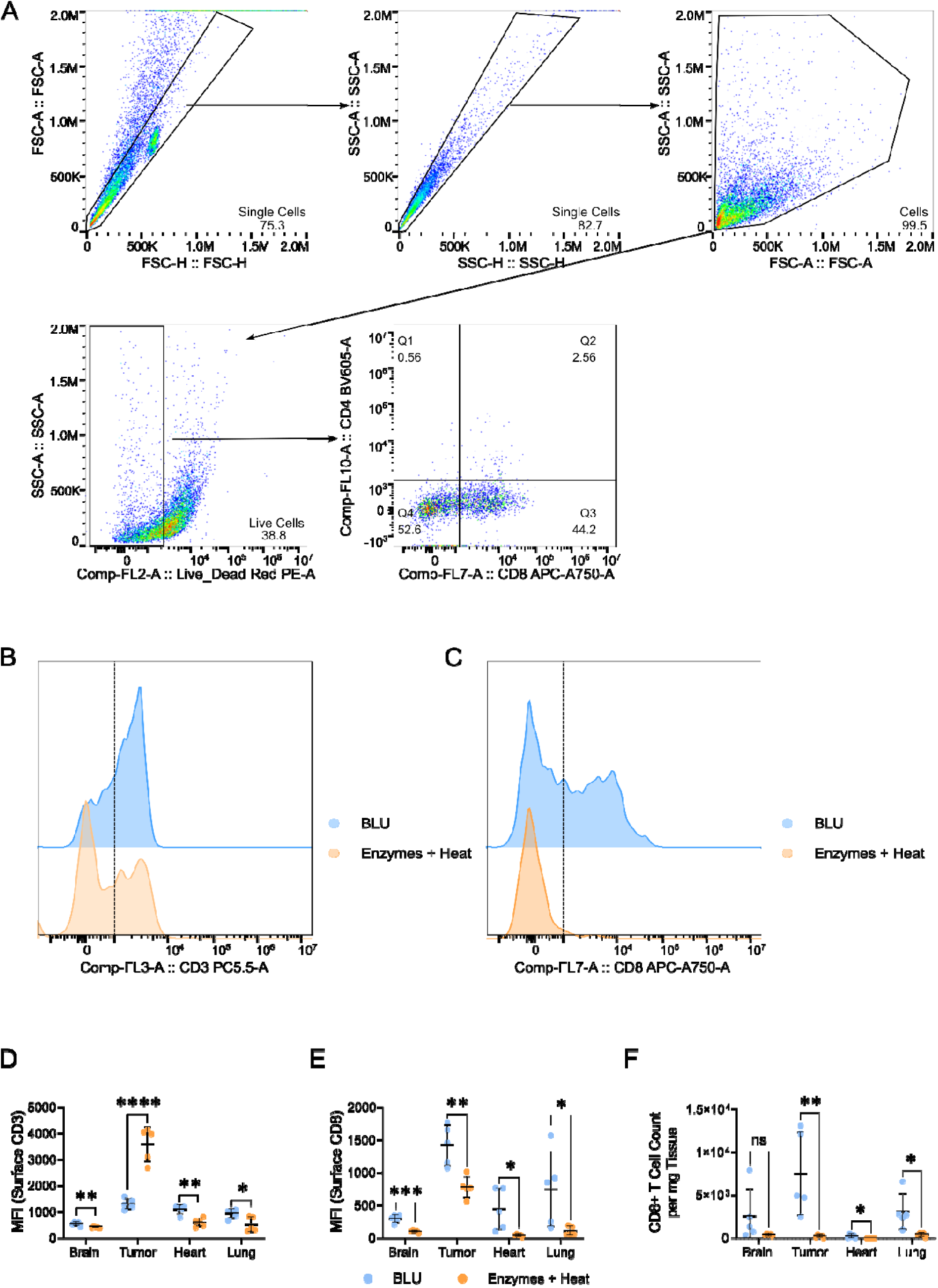
Significant differences in cell surface expression of T cell markers and CD8+ T cell counts were observed between enzymatic and BLU dissociations. (A) Representative gating strategy for identifying live, CD4+ and CD8+ cell populations (in heart tissue). Representative comparison of (B) CD3 and (C) CD8 expression, shown as mean fluorescence intensity (MFI) on all live cells, as determined by gating demonstrated above. (D) Quantification of CD3 cell surface expression and (E) CD8 cell surface expression on all live cells. (F) CD8+ T cell counts per milligram of tissue. ns = not significant, * = p-value ≤ 0.05, ** = p-value ≤ 0.01, *** = p-value ≤ 0.001, **** = p-value ≤ 0.0001.

For tissue types where very few or no detectable CD8+ T cells were isolated with enzyme and heat approaches, including brain, tumor and heart tissue (see Fig. 2F), further studies comparing expression markers on this cell subset were not possible. Wanting to validate the identity of those CD8+ T cells isolated by BLU, we assessed the expression of other known markers found on T cells, CD3 and CD45, (Supp. Fig. 1). For all tissue types, T cells identified as CD4-CD8+ by our gating strategy had increased expression of both CD45 and CD3 relative to the CD4-CD8-population.

### 3.3. Myeloid cell markers are diminished by enzymatic dissociation

Depending on the tissue type and the expected infiltrating immune cell populations, different myeloid cell panels were selected and evaluated (Supp. Tables 2-4). Despite minimizing variability in input material, several tissue types experienced significant loss of marker expression when dissociated with enzymes (Fig. 4). A representative gating strategy from tumor tissue is shown in Fig. 4A, where single cells were initially selected and a viability dye was used to exclude dead cells from further analysis.

**Figure 3.**
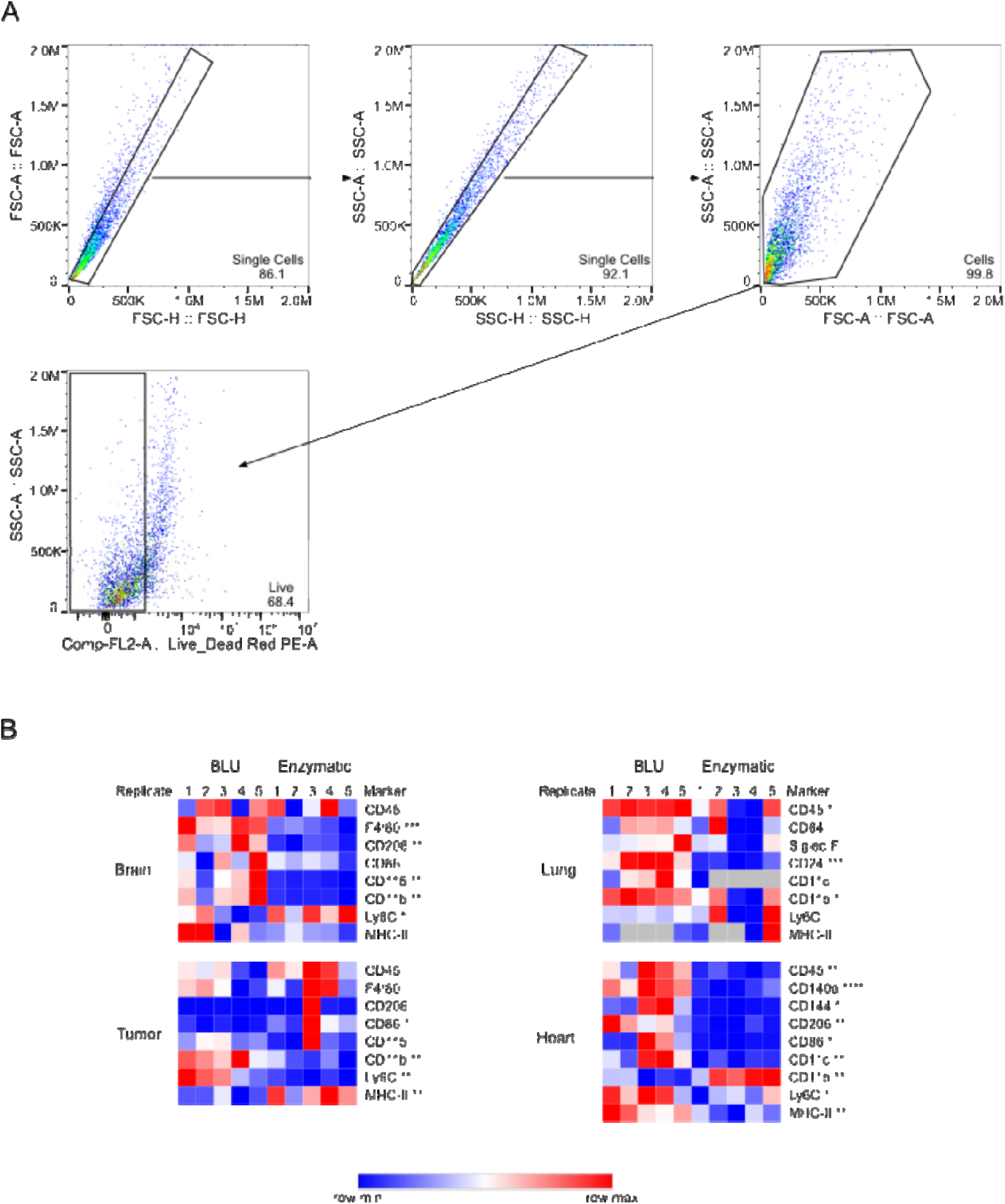
Marker expression on cells isolated from murine tumor, brain, lung, and heart tissue is significantly reduced by enzymatic dissociation. (A) Representative gating strategy for isolation of live cell populations using tumor tissue. (B) Comparison of median fluorescent intensity (MFI) signals of expression for several expected myeloid cell surface markers on live cells dissociated from different tissue types by either enzymes and heat or by BLU. Non-significant differences are not marked. * = p-value ≤ 0.05, ** = p-value ≤ 0.01, *** = p-value ≤ 0.001, **** = p-value ≤ 0.0001.

In brain tissue, these most notably consisted of microglia marker CD115, macrophage differentiating markers F4/80 and CD206, and dendritic cell marker CD11b (21,22). Monocyte marker Ly6C expression was significantly lower for cells dissociated by BLU. In tumor tissue, significant differentials were seen with both dissociation by BLU and by enzymes and heat, leading to inconclusive results. In heart tissue, expression of important mesenchymal and endothelial cell markers, CD140a and CD144, was significantly lost by enzymatic dissociation (23,24). Also in heart, expression of leukocyte marker CD45, macrophage differentiation marker CD206, and dendritic cell markers MHC-II, Ly6C, CD86, and CD11c was significantly lower in cells dissociated by enzymes and heat. Expression of the dendritic cell marker CD11b was increased by enzymatic dissociation relative to BLU, but it has previously been demonstrated that this marker is artificially upregulated during stress due to dissociation by enzymatic approaches (25). In lung, the leukocyte marker CD45, the broad eosinophil, neutrophil, and dendritic cell marker CD24, and the dendritic cell marker CD11b all had artificial losses of expression due to dissociation by enzymes and heat (26).

Across these four tissue types, artificial loss of expression was observed by enzymatic and heat-based dissociation on nearly all markers, suggesting a concern with cell subpopulation loss or cell surface expression interference of tissue dissociation by conventional dissociation methods using enzymes and heat.

## 4. Discussion

Tissue dissociation into single cell suspensions is an essential biological technique that enables a range of analyses key to inform downstream diagnostic development. Because tissue types and structures differ significantly throughout an organism, the conventional approaches to dissociating tissue are widely varied. The incorporation of heat or enzymes to enable this dissociation, however, is almost universal (27). The treatment of tissues with enzymes, heat, or a combination of the two can impact the resulting cell dissociations in ways that may influence conclusions drawn from surrounding research (12).

In this work, we have used flow cytometry to compare cell surface marker expression of immune cells dissociated from B16 melanoma tumors and from murine brain, heart, and lung tissue by either enzymes and heat or by BLU energy. While both methods yield comparable live cell counts per milligram of dissociated tissue and viability of single cell suspensions, we consistently see loss of cell marker expression for cells dissociated by enzymes and heat. On enzyme-dissociated cells from all tissue types besides tumor, CD3 expression was also significantly reduced. Even despite the lower expression of CD3 in single cell tumor suspensions dissociated by BLU, we observed significantly more CD8+ T cells in all suspensions aside from those dissociated from brain tissue. This corresponded with the significant loss of CD8 expression due to enzymatic dissociation, consistent with previous findings from Autengruber et al. (28).

When we examined other cell surface markers specific to myeloid lineages, we similarly observed significant losses in expression by enzyme dissociation. In brain tissue, CD115, F4/80, CD206, CD11b and Ly6C all had significantly lower expression in cells dissociated enzymatically. A subset of these markers, CD115, F4/80 and CD11b are broadly expressed markers on brain-based murine myeloid cells, used consistently for confirmation of cell subpopulations (21). Ly6C is generally expressed on inflammatory monocytes which infiltrate the brain pre-differentiation into microglia, and the presence of Ly6C^hi^ cells may be indicative of infection or illness (22). Similarly, CD206 expression corresponds to important border-associated murine macrophages, and information on this cell type was reduced by enzymatic dissociation (29). We were able to successfully observe the expected infection-induced immune activation in cells dissociated by BLU, but many of these signals were diminished or lost with enzymatic dissociation.

From the tumor, differential expression of signals was less conclusive. While dendritic cell marker CD11b and monocyte marker Ly6C expression were preserved by BLU dissociation, MHC-II and CD86 expression was higher in enzyme dissociated cells (11). While this may be due to the nature of differences in dissection of the tumor, a highly heterogenous tissue type, further work is needed to characterize this difference.

From the heart, multiple signals for different cell types were lost. Expression of CD140a, a mesenchymal marker, and CD144, an endothelial cell marker, were both significantly reduced in enzyme dissociated cells (23,24). Macrophage marker CD206 and dendritic cell markers MHC-II, Ly6C, CD86, and CD11c are all indicative of distinct immune subpopulations and were diminished or lost by enzymatic dissociation (26,30–32). The reduced expression of these markers in heart tissue by enzymatic dissociation yield an incomplete understanding of cardiac immune cell function.

Finally, in lung tissue, expression of CD45, CD24 and CD11b were significantly artificially reduced by enzymatic dissociation. CD45 and CD24 are broad cell markers expressed on multiple subpopulations of murine myeloid lineages in the lung commonly used for initial myeloid cell identification (33). CD11b is also a broadly expressed marker on murine dendritic cells that is often used to differentiate subpopulations (26). Reduced or lost expression of these markers leads to an incomplete understanding of circulating and tissue resident immune cell populations in lung tissue. However, additional repeats of lung tissue experiments are required to confirm these findings.

## 5. Conclusion

While our comparison of murine tissue dissociation measured stark differences in cell surface expression markers across BLU and enzymatic dissociation methods, further work is needed to determine whether the loss of these markers is limited to the cell surface or is indicative of the loss of entire cell populations. Findings of another study, where a comparison of an alternative mechanical dissociation method to the same commercially available enzymatic dissociation kits documented the loss of larger cell populations by enzymatic dissociation, suggest the latter (34). Regardless, our findings demonstrate that important immune cell markers are often compromised or lost by enzymatic dissociation, which may mislead conclusions about immune cell infiltration and localized immune responses. Our alternative, enzyme and heat-free acoustic dissociation by BLU energy, offers a method which preserves these markers, leading to an improved understanding of immune cell function from various tissue types. While BLU energy may not currently be optimized to dissociate all tissue types, as evidenced by the inconclusive findings with several expression markers in tumor, future work will focus on developing improved conditions of BLU energy to further enhance single cell suspensions of all tissue types and continue aiding researchers in their exploration of tissue landscapes and the corresponding role of immune cells.

## Author Contributions

M.A.M. analyzed and interpreted data and drafted the manuscript. M.B. ran experiments and edited the manuscript. S.Q. drafted the manuscript. B.Q. designed experiments and edited the manuscript. J.P. and S.A. assisted with running experiments. V.V. and S.L. developed instrumentation and conceptualized the project. K.R. oversaw data collection and analysis and drafted and edited the manuscript.

