## Supplemental Materials for "An enzyme-free, cold-process acoustic method for gentle and effective tissue dissociation"

**Supplemental Table 1.** Replicate tissue weights for tumor, brain, heart, and lung tissues.

| **Tissue Type** | **Dissociation Condition** | **Replicate** | **Weight (mg)** |
| --- | --- | --- | --- |
| **Tumor** | BLU | 1 | 84.3 |
|  |  | 2 | 102.7 |
|  |  | 3 | 99.4 |
|  |  | 4 | 100.6 |
|  |  | 5 | 101.5 |
|  | Enzymes and Heat | 1 | 95.9 |
|  |  | 2 | 88.2 |
|  |  | 3 | 100.6 |
|  |  | 4 | 87.5 |
|  |  | 5 | 83.8 |
| **Brain** | BLU | 1 | 120.0 |
|  |  | 2 | 113.4 |
|  |  | 3 | 114.8 |
|  |  | 4 | 121.3 |
|  |  | 5 | 116.2 |
|  | Enzymes and Heat | 1 | 112.8 |
|  |  | 2 | 124.0 |
|  |  | 3 | 115.8 |
|  |  | 4 | 118.0 |
|  |  | 5 | 113.7 |
| **Heart** | BLU | 1 | 106.8 |
|  |  | 2 | 111.3 |
|  |  | 3 | 109.1 |
|  |  | 4 | 109.8 |
|  |  | 5 | 107.7 |
|  | Enzymes and Heat | 1 | 121.7 |
|  |  | 2 | 129.1 |
|  |  | 3 | 105.3 |
|  |  | 4 | 122.5 |
|  |  | 5 | 102.3 |
| **Lung** | BLU | 1 | 95.1 |
|  |  | 2 | 109.9 |
|  |  | 3 | 114.0 |
|  |  | 4 | 96.3 |
|  |  | 5 | 115.5 |
|  | Enzymes and Heat | 1 | 100.4 |
|  |  | 2 | 103.7 |
|  |  | 3 | 97.6 |
|  |  | 4 | 107.7 |
|  |  | 5 | 100.3 |

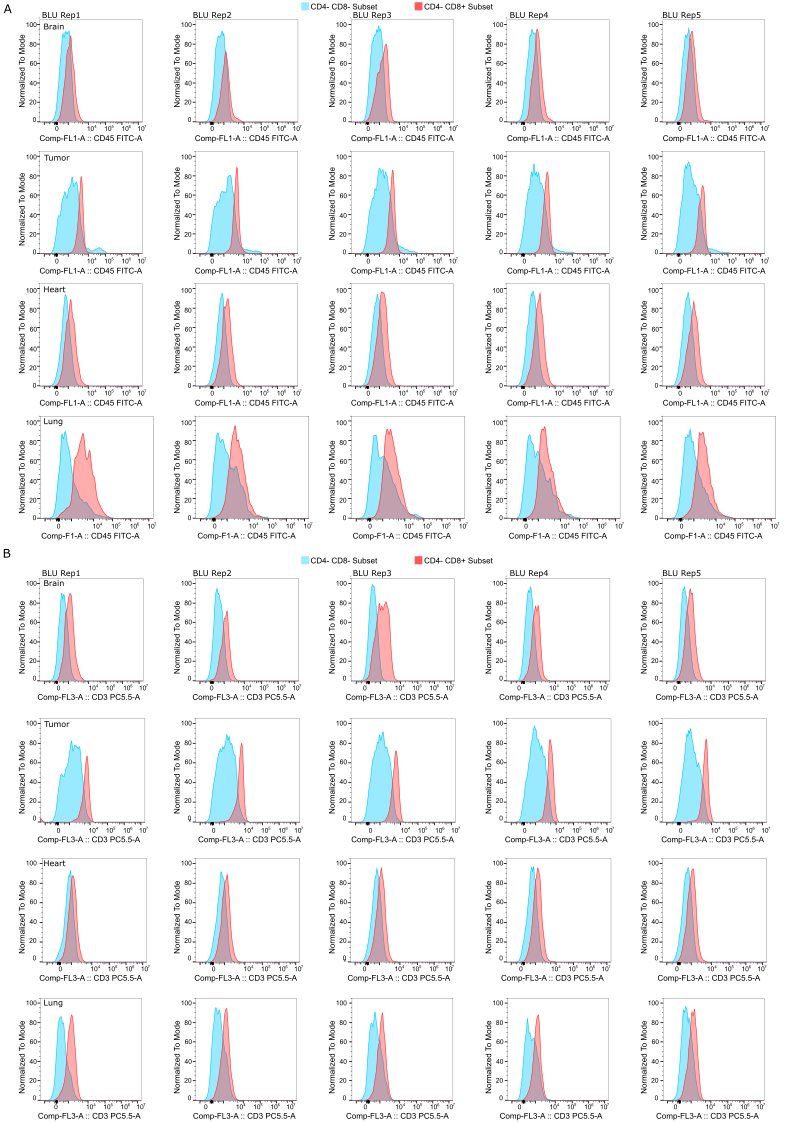

**Supplemental Figure 1.** Cell surface co-expression of common T cell markers on subpopulations of BLU™ dissociated tissue samples identified as CD4-CD8+ relative to the subpopulation identified as CD4-CD8-. (A) Co-expression of CD45. (B) Co-expression of CD3.

**Supplemental Table 2.** Myeloid panel for monocytes and macrophages found in tumor and brain tissues.

| **Cell Surface Marker** | **Fluorochrome** | **Manufacturer** | **Catalog #** | **Dilution** |
| --- | --- | --- | --- | --- |
| CD115 | AF750 | Biolegend | 135536 | 1:400 |
| CD11b | BV421 | Biolegend | 101251 | 1:400 |
| Ly6C | BV510 | Biolegend | 128033 | 1:400 |
| MHC-II | BV605 | Biolegend | 107639 | 1:400 |
| F4/80 | PE-Cy7 | Biolegend | 139314 | 1:400 |
| CD86 | AF700 | Biolegend | 123130 | 1:500 |
| CD206 | AF647 | Biolegend | 151508 | 1:500 |
| CD45 | FITC | Biolegend | 103108 | 1:500 |
| Live/Dead Red | PE | ThermoFisher | L34971 | 1 µL/mL |

**Supplemental Table 3.** Heart-specific myeloid panel with markers for monocytes and macrophages found in heart tissues.

| **Cell Surface Marker** | **Fluorochrome** | **Manufacturer** | **Catalog #** | **Dilution** |
| --- | --- | --- | --- | --- |
| CD11c | APC/Fire750 | Biolegend | 117351 | 1:400 |
| CD11b | BV421 | Biolegend | 101235 | 1:400 |
| Ly6C | BV510 | Biolegend | 128033 | 1:400 |
| MHC-II | BV605 | Biolegend | 107639 | 1:400 |
| CD144 | PE-Cy7 | Biolegend | 138015 | 1:400 |
| CD140a | PerCp-Cy5.5 | Biolegend | 135913 | 1:400 |
| CD86 | AF700 | Biolegend | 105023 | 1:500 |
| CD206 | AF647 | Biolegend | 103123 | 1:500 |
| CD45 | FITC | Biolegend | 103108 | 1:500 |
| Live/Dead Red | PE | Invitrogen | L34972 | 1 µL/mL |

**Supplemental Table 4.** Lung-specific myeloid panel with markers for monocytes and macrophages found in lung tissues.

| Cell Surface Marker | Fluorochrome | Manufacturer | Catalog # | Dilution |
| --- | --- | --- | --- | --- |
| CD11c | APC/Fire750 | Biolegend | 117351 | 1:400 |
| CD11b | BV421 | Biolegend | 101235 | 1:400 |
| Ly6C | BV510 | Biolegend | 128033 | 1:400 |
| MHC-II | BV605 | Biolegend | 107639 | 1:400 |
| CD64 | PE-Cy7 | Biolegend | 161007 | 1:400 |
| CD24 | AF700 | Biolegend | 101835 | 1:500 |
| Siglec F | AF647 | Biolegend | 155519 | 1:500 |
| CD45 | FITC | Biolegend | 103108 | 1:500 |
| Live/Dead Red | PE | Invitrogen | L34972 | 1 µL/mL |
